# *Chironomus riparius* (Diptera) genome sequencing reveals the impact of minisatellite transposable elements on population divergence

**DOI:** 10.1101/080721

**Authors:** Ann-Marie Oppold, Hanno Schmidt, Marcel Rose, Sören Lukas Hellmann, Florian Dolze, Fabian Ripp, Bettina Weich, Urs Schmidt-Ott, Erwin Schmidt, Robert Kofler, Thomas Hankeln, Markus Pfenninger

## Abstract

Active transposable elements (TEs) may result in divergent genomic insertion and abundance patterns among conspecific populations. Upon secondary contact, such divergent genetic backgrounds can theoretically give rise to classical Dobzhansky-Muller incompatibilities (DMI), a way how TEs can contribute to the evolution of endogenous genetic barriers and eventually population divergence. We investigated whether differential TE activity created endogenous selection pressures among conspecific populations of the non-biting midge *Chironomus riparius,* focussing on a *Chironomus*-specific TE, the minisatellite-like *Cla-element*, whose activity is associated with speciation in the genus. Using an improved and annotated draft genome for a genomic study with five natural *C. riparius* populations, we found highly population-specific TE insertion patterns with many private insertions. A highly significant correlation of pairwise population F_ST_ from genome-wide SNPs with the F_ST_ estimated from TEs suggests drift as the major force driving TE population differentiation. However, the significantly higher *Cla-element* F_ST_ level due to a high proportion of differentially fixed *Cla-element* insertions indicates that segregating, i.e. heterozygous insertions are selected against. With reciprocal crossing experiments and fluorescent in-situ hybridisation of *Cla-elements* to polytene chromosomes, we documented phenotypic effects on female fertility and chromosomal mispairings that might be linked to DMI in hybrids. We propose that the inferred negative selection on heterozygous *Cla-element* insertions causes endogenous genetic barriers and therefore acts as DMI among *C. riparius* populations. The intrinsic genomic turnover exerted by TEs, thus, may have a direct impact on population divergence that is operationally different from drift and local adaptation.

## Introduction

It is common knowledge that the divergence of geographically isolated populations is shaped by genomic drift and local adaptation. However, less attention has been given to the contribution of structural genetic elements (e.g. transposable elements (TEs), satellite repeats, duplications; Bierne *et al.* 2011; Seehausen *et al.* 2014) to population divergence. Structural variation of genomes in different populations, as a result of drift and intrinsic genomic interactions, may give rise to endogenous incompatibilities upon secondary contact of diverged populations (Bierne *et al.* 2011). Endogenous incompatibilities due to structural genomic variation are thus just a special case of the more general concept of Dobzhansky-Muller incompatibilities (DMI) that can arise from any sort of interacting loci or any functionally relevant mutation even between conspecific populations of otherwise weakly differentiated genetic backgrounds (Barton & de Cara 2009; Orr & Turelli 2001). They can therefore contribute to the evolution of reproductive isolation (RI) and ultimately speciation (Dover 1982).

Comprising a considerable fraction of most eukaryotic genomes, especially TEs have been proposed as major drivers of genome evolution and were suggested to play an important role in events leading to speciation (Kazazian 2004; Werren 2011). TEs evolve rapidly, transpose throughout the genome and may lead to chromosomal rearrangements by ectopic recombination (Jurka *et al.* 2011; Montgomery *et al.* 1991). Therefore, it is reasonable to consider TEs also as major drivers of genomic diversification among populations of the same species. In general, TE insertions in the genome are deleterious (Pasyukova *et al.* 2004) and therefore mechanisms for the suppression of TE spread are prevalent positively selected (Simkin *et al.* 2013). If different TE suppression mechanisms evolved in two populations, unstable genome constellations or uncontrolled TE spread in hybrids and eventually reduced hybrid fitness can be the consequence (Crespi & Nosil 2013). However, besides the sensitive balance between TE proliferation and its suppression by the host genome, divergent insertion patterns of TEs and other genetic elements might also lead to other forms of DMI in hybrid individuals of different populations (Bierne *et al.* 2011), and hybridization among divergent populations may thus provoke substantial genomic stress (Fontdevila 1992).

Empirical evidence for the potential involvement of TEs in postzygotic RI comes e.g. from *Drosophila,* where three different TEs (P-, I-, and hobo element;Kavi *et al.* 2005; Kofler *et al.* 2015; Simmons *et al.* 2015) are responsible for hybrid dysgenesis, which involves elevated rates of mutation, chromosomal rearrangements, illicit recombination in males or sterility of females due to gonadal dysfunction (Bingham *et al.* 1982; Bucheton *et al.* 1984; Galindo *et al.* 1995). Similar TE-mediated hybrid breakdown phenomena have been observed in plants (Martienssen 2010) and vertebrates (Dion-Cote *et al.* 2014). While these documented effects are due to incompatibilities of classical transposase-encoding TE types, the role of non-classical TEs, which can be organized as tandemly repeated, interspersed minisatellite clusters, in the causation of DMI and hence RI remains unexplored. Such mobile repeat clusters could, however, have substantial impact, since they are often associated with chromosomal heterochromatin, thereby possibly affecting genomic stability (Ilkova *et al.* 2013), chromosome pairing (Hernandez-Hernandez *et al.* 2008), recombination (Talbert & Henikoff 2010), and expression of neighbouring genes by epigenetic position effects (Pezer & Ugarkovic 2012). During cell division, improper chromatid pairing may eventually lead to developmental problems and reduced hybrid fitness (Hailu *et al.* 1999; Oleszczuk & Lukaszewski 2014).

TEs have been invoked in the speciation of the morphologically cryptic non-biting midge sister species *Chironomus riparius* and *C. piger* (formerly called *C. thummi thummi and C. t. piger*). Depending on the crossing direction, two incompatibility syndromes involving hybrid dysgenesis have been described in these closely related midges, the Rud and the HLE syndrome (Hägele 1984, 1985, 1987, 1995; Hägele & Kasper-Sonnenberg 2000; Hägele & Lachmann 1992; Hägele *et al.* 1995; Hägele & Oschmann 1987). The latter also occurs on an intra-specific level in different *C. riparius* populations as aberrant traits such as reduced egg hatch and chromosome aberrations occurring with population-specific intensity (Hägele 1995). The molecular basis of the syndromes, however, has so far not been investigated.

On the genome level, a markedly different genome size and organization of repetitive DNA is suspected to contribute to the above syndromes (Schmidt 1984). The *C. riparius* genome is about 27% larger compared to *C. piger*, the latter presumably representing the ancestral chromosomal karyotype (Keyl 1965). One-fifth of the accumulated DNA in *C. riparius* is due to the transposable *Cla-element* sequence (Schmidt 1981). *Cla-elements* occur either as tandem-repetitive minisatellite-like clusters often containing more than 20 repeat units (Hankeln *et al.* 1994), or as monomeric repeats that share characteristics with SINE (short interspersed element) retrotransposons (Hankeln & Schmidt 1987). The exact mechanism of transposition of these sequences is yet unclear, and it is unknown whether *Cla-elements* are still active. In *C. piger* and the more distantly related *C. luridus*, *Cla-element* clusters are mostly restricted to centromeric regions, whereas in *C. riparius,* they additionally infest most chromosomal arms (Hankeln *et al.* 1994; Hankeln & Schmidt 1987). The AT-rich 120 bp consensus sequence of the *Cla-element* has the potential of forming hairpin structures and induces a strong sequence-directed DNA curvature, multimeric *Cla-elements* in particular (Hankeln 1990; Israelewski 1983; Schmidt 1981, 1984). Such structures may be particularly relevant with regard to potential effects on chromosome integrity, centromere formation and synapsis of homologous chromosomes. Comparable sequence-inherent curvature of the mouse major centromeric satellite DNA seems important for heterochromatin formation (Radic *et al.* 1987). DNA hairpins are involved in several biological processes and may affect transcription, recombination, and replication in pro- and eukaryotes (Bikard *et al.* 2010; Mariappan *et al.* 1996), as well as kinetochore formation in human centromers (Catasti *et al.* 1994). In *C. riparius*, common chromosomal breakpoints were reported to co-localise with heterochromatic regions that contain blocks of tandem-repetitive DNA such as the *Cla-element* (Bovero *et al.* 2002). As the *Cla-element* potentially affects chromosomal architecture, function and genome evolution in *C. riparius*, its genomic spread might hypothetically be involved in the observed genomic incompatibilities to its sister species, and therefore, be causal for speciation events in the genus.

We thus aimed at elucidating whether diverging patterns of *Cla-element* distribution contribute disproportionately to population divergence and may result in DMI between populations by creating endogenous genetic barriers in *C. riparius*. To this end, we sequenced, assembled, and annotated a *C. riparius* draft genome. In a genome-wide population comparison we investigated the population differentiation at SNP loci and at *Cla-element* insertion sites to see (i) if the *Cla-element* distribution and frequency is more diverged than expected from classical SNPs between populations. We then hybridised geographically distant *C. riparius* field populations, and by *in-situ* hybridisation of *Cla-element* clusters to polytene chromosomes we investigated (ii) whether interpopulation hybrid individuals showed phenotypic effects on the level of chromosomes as an evidence for DMI. With laboratory experiments we tested (iii) whether we find evidence for reduced hybrid fitness.

## Material and Methods

### Sequencing, assembly, scaffolding, and annotation of a *C. riparius* draft genome

Approximately 50 larvae of a *C. riparius* laboratory culture established several decades ago were used for extraction of genomic DNA. After quality checks of DNA, three paired-end and two mate-pair sequencing libraries of 3 kb and 5.5 kb insert size were prepared (Illumina, CA, USA) and run on Illumina HiSeq2500 and MiSeq instruments (Institute of Molecular Genetics, University of Mainz, Germany; StarSEQ, Mainz, Germany). Sequence reads were quality checked, quality processed and then assembled using the Platanus v1.2.1 pipeline (Kajitani *et al.* 2014) with kmer-sizes between k=32 and k=84. Iterative scaffolding of contigs was conducted by Platanus with default parameters and SSPACE 3.0 (Boetzer *et al.* 2011), the contig-extension option enabled. Completeness and integrity of the draft genome was examined by back-mapping of reads with BWA v0.7.10-r789 (Li & Durbin 2009) using the *bwa mem* algorithm, and by analyses using the tools BUSCO v1.1b1 (Simao *et al.* 2015) and REAPR 1.0.18 (Hunt *et al.* 2013).

The genome sequence was then annotated using three iterations of the MAKER2 v2.31.8 pipeline (Cantarel *et al.* 2008; Holt & Yandell 2011). For annotation of protein-coding genes we supplied the pipeline with a reference transcriptome assembled from cDNA sequence data of embryonic, larval and adult *C. riparius* specimen and additionally with the Swiss-Prot database (obtained on 2016-01-13 from uniprot.org). Repeat detection was based on a custom repeat library. See Supplementary Methods S1 for more information on the whole process of generating the “CRIP_Laufer” draft genome sequence and annotation. The reads are available at the Short Read Archive (SRA) with accession numbers XX-XX, the scaffolds are available from XXX. The annotation file as well as access to a *C. riparius* genome browser is available upon request.

### Sampling of natural populations

We sampled a total of five natural *C. riparius* populations from across Europe (Supplementary Table S1). Larvae were collected alive to establish laboratory cultures, as well as conserved in ethanol for pooled sequencing. Species identification was done with COI barcoding (Pfenninger *et al.* 2007) of F_1_clutches from the laboratory culture and all preserved larvae. To rule out the misidentification of possible hybrid individuals, a fragment of the nuclear-encoded mitochondrial 39S ribosomal protein L44 was added as nuclear marker. Barcoding of the two markers was performed as described in (Oppold *et al.* 2016).

### Pooled sequencing of natural populations

The Pool-Seq approach allows to obtain unbiased estimates of allele frequencies in the entire genome by sequencing a population pool of many (> 100) diploid individuals to relatively low coverage, thus making probable that each chromosome in the pool is sequenced only once per locus (Futschik & Schlotterer 2010). We obtained population genomic data for each of five natural *C. riparius* populations (Supplementary Table S1). We dissected head capsules from all barcoded larvae of the respective populations, pooled them and extracted the genomic DNA using the DNeasy Blood & Tissue Kit (QIAGEN, Hilden, Germany). DNA concentration was measured with the Qubit^®^ dsDNA BR Assay Kit in a Qubit^®^ fluorometer and quality was assessed by gel-electrophoresis. Sequencing libraries of the individual pools were constructed using the TrueSeq DNA Nano Library Prep Kit (Illumina, CA, USA) tagged with different multiplex barcodes (GENterprise Genomics, Mainz, Germany). For each library the same insert size was selected, ranging from 180-680 (average 380). All pools were sequenced as 100 bp paired-end sequences on an Illumina HiSeq 2500 platform (Institute of Molecular Genetics, University of Mainz, Germany). Adapter clipping and end-quality trimming on the raw sequences was performed with TRIMMOMATIC (ILLUMINACLIP:adapters.fa:2:30:10:8, Bolger *et al.* 2014) using a sliding-window approach (window size 4, Phred quality 20). Trimmed reads were inspected with FASTQC (v0.11.2; http://www.bioinformatics.babraham.ac.uk/projects/fastqc/).

### Inference of genome-wide population differentiation and heterozygosity

To estimate the genome-wide population differentiation and heterozygosity based on single nucleotide polymorphisms (SNPs), we followed the PoPoolation2 pipeline (Kofler *et al.* 2011b). The Pool-Seq reads of the five natural populations were mapped to the draft genome with BWA using the *bwa aln* algorithm with a maximum number of 1 gap per alignment, and a mismatch probability of 0.01. Additionally, seeding was disabled, as recommended by (Kofler *et al.* 2011a). Paired-end information was linked with the *bwa sampe* algorithm. Mapping success was inspected with QualiMap v2.0 (Okonechnikov *et al.* 2016). Downstream processing of the bam files strictly followed the pipeline of PoPoolation2 (Kofler *et al.* 2011b). Mean coverage was calculated with R and *samtools depth* (SAMtools utilities; version 1.1, (Li *et al.* 2009). To ensure comparable mean coverage between the data, the bam-file of the Lorraine population (NMF) was downsampled to the average coverage of the population-means with Picard (DownsampleSam.jar; v1.119, available at http://picard.sourceforge.net). SNP-calling was performed on the sync-file that combined all five Pool-Seq data sets. To calculate population-specific heterozygosity, we called SNPs with the *snp-frequency-diff.pl* script of the PoPoolation2 package (Kofler *et al.* 2011b) with a minimum count of 4, and a coverage ranging from a minimum of 20 to the upper 2% of the respective data set (i.e. MF: 50, MG: 37, NMF: 49, SI: 50, SS: 45). From the output (rc-file) we calculated allele frequencies and heterozygosity for all population-specific SNPs, as well as for population private SNPs. To estimate the pairwise F_ST_ for each SNP between all possible population pairs, we used the *fst-sliding.pl* script of the PoPoolation2 package (parameters: --min-count 4 --min-coverage 20 --max-coverage 50,37,67,50,45 --min-covered-fraction 1.0 --window-size 1 --step-size 1 --pool-size 168:112:105:155:118 --suppress-noninformative). Mean heterozygosity and mean F_ST_s for each pairwise comparison were then calculated in R.

### Estimating TE abundance with PoPoolationTE

As required for PoPoolationTE, we combined the *C. riparius* draft genome and a custom library of 108 *Chironomus*-specific transposable elements and repeats, including the 117 bp *Cla-element* consensus sequence (Schmidt 1984). To test whether all *Cla-element* variants present in our data can be mapped to the *Cla-element* consensus sequence, we first performed a BLAST (Altschul *et al.* 1990) search of all available *Cla-element* sequences from NCBI (gi|6456818, gi|7099, gi|89994172, gi|89994177, gi|89994178, gi|89994179) against the five Pool-Seq data sets (options: -outfmt 6-max_target_seqs 100000000 -evalue 1e-30). The output of each population and query was filtered for a minimum alignment length of 80 bp. We then merged the filtered output that contained a single BLAST hit per population to at least one query (filtering duplicates with a custom perl script). This resulted in a file containing all the *Cla-element* variation present in our five data sets. 98.4% of this data could successfully be back-mapped (*bwa bwasw*, see below) to the consensus sequence of the *Cla-element*, which demonstrated that this sequence is a suitable reference for our study.

The custom TE-library was first used for masking the genome with RepeatMasker v4.0.5 (Smit *et al.* 2013-2015). The masked genome was then combined with the TE-library and further used as combined reference sequence for the mapping of the five Pool-Seq data sets. We generated a TE-hierarchy, which is required for PoPoolationTE by obtaining the insert name, order, family, and subfamily for every entry in the TE-library. Paired-end sequence reads of the five Pool-Seq data sets were mapped as singletons to the combined reference sequence with BWAs Smith-Waterman alignment algorithm *bwasw* (Li & Durbin 2010). Paired-end information was *post hoc* recovered with the samro.pl script from the PoPoolationTE tool package v51 (Kofler *et al.* 2012). The resulting paired-end bam files were subsampled to an equal number of 50 million reads using *samtools view* v1.1, (Li *et al.* 2009), in order to avoid biased estimations of TE-insertions when comparing the different populations. Next, we sorted the sam files (*samtools sort*) of the five populations and identified TE-insertions in each sample separately (and the population frequency of the insertions) with PoPoolationTE (Kofler *et al.* 2012). Based on the annotation file, we investigated whether TE-insertions were localised close to coding regions, in introns or exons (custom script).

### Population genomic analyses of *Cla-element* insertions

The genomic localization of TE-insertions estimated by PoPoolationTE may be slightly inaccurate and thus result in a slight variation of the genomic position of identical TE insertions in different Pool-Seq data sets. In order to compare TE abundance between samples, we clustered TE insertion sites from the five natural populations within a specific genomic window to one single TE-insertion using a custom script. For the identification of the optimal window-size, we calculated the exponential one-phase decay model of increasing window-size with the number of TE-insertions in GraphPad Prism^®^ (Version 5, GraphPad Software, San Diego, USA; Supplementary Figure S1). A plateau was reached at a window size of about 450 bp for a model using all TEs excluding the *Cla-element*, as well as for the model using solely the *Cla-element*. All downstream analyses were therefore based on homogenisation of TE-insertions within a 450 bp window across populations.

The *Cla-element* is a tandem-repetitive element, building large clusters in the genome of *C. riparius* (Hankeln *et al.* 1994). With PoPoolationTE it is, however, not possible to distinguish monomer insertions from clusters (Kofler *et al.* 2012). To quantify the mean *Cla-element* cluster size we first predicted an expected number of reads per monomer insertion by combining the number of identified insertion sites with the fraction of reads that cover one monomer and the genome-wide mean coverage adjusted to the mean frequency (Table 1, calculation steps outlined in column 1). We extracted the reads mapping to the *Cla-element* directly from the reads mapped to the TE-combined-reference (using *samtools view*). Comparison of observed to expected values finally gave an average cluster size estimate. We randomly selected low-frequency *Cla-element* insertions from the PoPoolationTE output and validated them visually with the Integrative Genomics Viewer v2.3.68 (Robinson *et al.* 2011; Thorvaldsdottir *et al.* 2013).

**Table 1:** Estimation of *Cla-element* cluster size in the genome of different *C. riparius* populations (MF: Rhone-Alpes, MG: Hessen, NMF: Lorraine, SI: Piemont, SS: Andalusia).

|  |  | MF | MG | NMF | SI | SS |
| --- | --- | --- | --- | --- | --- | --- |
| **A** | insertion sites | 849 | 538 | 807 | 803 | 772 |
| **B** | mapped reads to *Cla* (**observed value**) | 1918365 | 1494769 | 2117883 | 1761580 | 1889805 |
| **C** | mean read length | 79.0 | 75.5 | 75.5 | 79.0 | 75.5 |
| **D** | fraction of monomer coverage by one read | 1.48 | 1.55 | 1.55 | 1.48 | 1.55 |
| **E** | genome-wide mean coverage | 24 | 23 | 22 | 24 | 23 |
| **F** | mean *Cla* frequency | 0.84 | 0.87 | 0.84 | 0.84 | 0.82 |
| **G=**<br>**A\*D\*E\*F** | predicted number of reads per monomer (**expected value**) | 25390 | 16710 | 23163 | 24078 | 22476 |
| **H=B/G** | predicted mean cluster size (**observed/expected**) | 75.6 | 89.5 | 91.4 | 73.2 | 84.1 |
| **I=**<br>**B/(D\*E\*F)** | estimated genome-wide copy number | 64147.2 | 48126.3 | 73786.1 | 58748.5 | 64909.4 |

As an additional line of evidence for potential ongoing activity of the *Cla-element*, we scanned the publicly available transcriptome datasets SRR834592 and SRR834593 (Schmidt *et al.* 2013) for *Cla-element*-containing transcripts by mapping those reads to the consensus sequence of the *Cla-element* dimer with the *bwa bwasw* algorithm.

To visualise the overlap of *Cla-element* insertions between the five populations, we plotted Venn diagrams on the basis of all *Cla-element* insertions identified with PoPoolationTE (online-tool: http://bioinformatics.psb.ugent.be/webtools/Venn/ accessed 29.01.2016), for all and for fixed insertions. With the relation of private *Cla-element* insertions and the total number per population we performed population-pairwise comparisons with Chi² tests. The private *Cla-element* insertions were further related to the annual mean temperature of the respective natural population site (climate data was derived from the WorldClim database; Hijmans *et al.* 2005) as a non-linear exponential one-phase association in GraphPad Prism^®^.

Based on the frequency estimates of TE-insertions, we calculated expected heterozygosity and pairwise F_ST_ for all TEs and solely for *Cla-element* insertions. Expected heterozygosity was calculated for private insertions in each population. F_ST_ was calculated as the reduction in heterozygosity in subpopulations (H_S_) compared to heterozygosity of the total population (H_T_): F_ST_ = (H_T_-H_S_)/H_T_. Isolation-by-distance (IBD) was analysed by correlating the pairwise F_ST_ with the geographical distance between the sampling sites of populations. Statistical significance of the correlations was assessed with Pearson’s *r* and a two-tailed *p*-value for the F_ST_ correlation and a one-tailed *p*-value for the IBD pattern. Mean heterozygosity and mean pairwise F_ST_ were then calculated in R. The mean heterozygosity of private *Cla-element*s and TEs over all populations was compared to genome-wide heterozygosity calculated for SNPs. For all populations, we compared the proportion of fixed insertions private to a population with fixed private SNPs. Statistical significance was tested using one-way ANOVA with Bonferroni correction.

To infer whether potential endogenous selection on TEs affected levels of population divergence at SNPs in their vicinity, we compared the mean F_ST_ of SNPs within 50, 100 and 500 bp upstream and downstream of *Cla-elements* and other TE insertion sites with the F_ST_s from 10,000 randomly chosen SNPs from other locations in the genome, respectively.

### Hybridisation experiments

Barcoded egg clutches of the natural populations were used to start pure *C. riparius* laboratory cultures that were acclimatized to standard conditions for at least three generations. Hybridisation experiments were performed with the three geographically most distant populations (Hessen, Piemonte, Andalucia) to test for the degree of intraspecific RI. The experiment comprised two subsequent generations (F_1_-F_2_), whereas the hybridisation happened during the adult stage of F_1_. We hybridized the populations in both crossing directions, screening the F_2_ generation for patterns of RI by following the life-trait parameters: mortality, mean emergence time (EmT_50_), clutches per female, eggs per clutch, fertility of clutches, and population growth rate (PGR). Different groups were compared statistically via one-way ANOVA with Tukey post-test (mortality, EmT_50_, PGR) in GraphPad Prism^®^. Population-specific values (clutches per female, fraction of fertile clutches) were compared as contingency tables with Fisher’s exact test in R v3.2.3, (R-Core 2015). For each population, 360 first-stage larvae (L1) were distributed over six replicates and kept under constant condition at 20°C (Oppold *et al.* 2016). Larvae were fed with a developmental stage-adjusted amount of ground TetraMin (Tetra GmbH, Germany). After emergence of F_1_ imagines, 50 individuals of each sex were distributed to separate reproduction cages to start hybridisation populations as well as pure control populations. Egg clutches were removed daily from the reproduction cages and stored separately in 6-well plates with medium under the same conditions to document the early development of each clutch and define its fertility with the successful hatch of larvae as criteria. After hatch, all larvae of a respective pure population or hybridisation population were pooled to start the F_2_ generation. 40 larvae were distributed over five replicates during larval development (total of 200 larvae per population) and after emergence 50 individuals of each sex from a respective pure or hybridisation population were distributed to reproduction cages.

### Fluorescent in-situ hybridisation of *Cla-elements*

Salivary gland polytene chromosomes of *C. riparius* were dissected from salivary glands of L4 larvae from the F_2_ generation of the hybridisation experiment, i.e. the pure populations as well as hybrids from all crossing directions. The preparation of chromosomes and *in-situ* hybridisation of *Cla-elements* followed (Hankeln & Schmidt 1987; Schmidt *et al.* 1988). As *Cla-element* probe we used clone pAD 2903 containing a *Cla-element* dimer (Hankeln 1990), which was DIG-labelled (DNA Labeling Kit^©^, Roche Life Science, USA). FITC-conjugated anti-DIG (Roche Life Science, USA) was used as antibody to detect hybridization sites. Chromosomes were DAPI-stained (Roche Life Science, USA). Preparations were inspected with a fluorescence microscope system (Olympus BX61 + Olympus BX UCB, Olympus, Japan). Pictures of DAPI-stained chromosomes were overlaid by pictures of the *Cla-element* FISH in Photoshop CS6 Extended (Adobe, USA).

## Results

### *C. riparius* draft genome

We generated about 350 million Illumina sequence reads from five sequencing libraries. On average 71.5 % of the reads passed quality filtering steps, resulting in a total 250 million reads and approximately 130x genome coverage (Supplementary Table S2). The assembly resulted in 41,974 contiguous sequences ≥ 500 bp with a GC content of 31%. These contigs could be interconnected to yield 5,292 scaffolds ≥ 1,000 bp with an N50 value of 272,065 bp (NG50 = 227,750). The total length of the draft genome is 180,652,019 bp (16 % N’s, Supplementary Table S3), which covers about 90 % of the estimated genome size of 200 Mb (Schmidt-Ott *et al.* 2009). On average, 91.9% of the reads used for assembly could successfully be mapped back against the draft genome (Supplementary Table S4). The draft genome contains 93 % of a set of 2,675 core arthropod genes (Supplementary Table S5). Using read pairing information, 550 (0.1%) scaffolding positions were identified that cannot be resolved unambiguously.

The genome was annotated using a reference transcriptome assembled from 14 cDNA datasets (Supplementary Table S6). We found predictions for 13,093 protein-coding genes with on average 4.7 exons per gene. The exons have an average length of 355 bp and all exons together sum up to 11% of the whole genome (Supplementary Table S7). Additionally, 71,990 regions were marked as repetitive element sequences, including 582 potential *Cla-element* positions.

### Transposable element families in *C. riparius*

We identified 19 different TE families summing up to an overall number of 5,645 insertion sites within all five natural *C. riparius* populations (Fig. 1). These TE families are all present at different insertion numbers in the five populations. Alu, Hind, and *Cla* minisatellite-like families, as well as the CTRT1 SINE element, the NLRCth1 retrotransposon and the TECth1 DNA transposon each occurred at more than 100 insertion sites across the genome in all populations (Fig. 1). The majority of TE insertion sites were fixed within populations (Supplementary Figure S3). However, in all populations we also found low to intermediate insertion frequencies for all TE families, except for the coding tandem repeats originating from Balbiani Ring genes. Partial detection of a subtelomeric repeat known from *C. dilutus* (gi|6525125 - gi|6525127) could not be localized in the draft genome but was found to be associated to the *C. thummi* DNA for TsB and TsC telomeric tandem repeat in the repeat library (gi|9557882 and gi|9557883).

**Figure 1:**
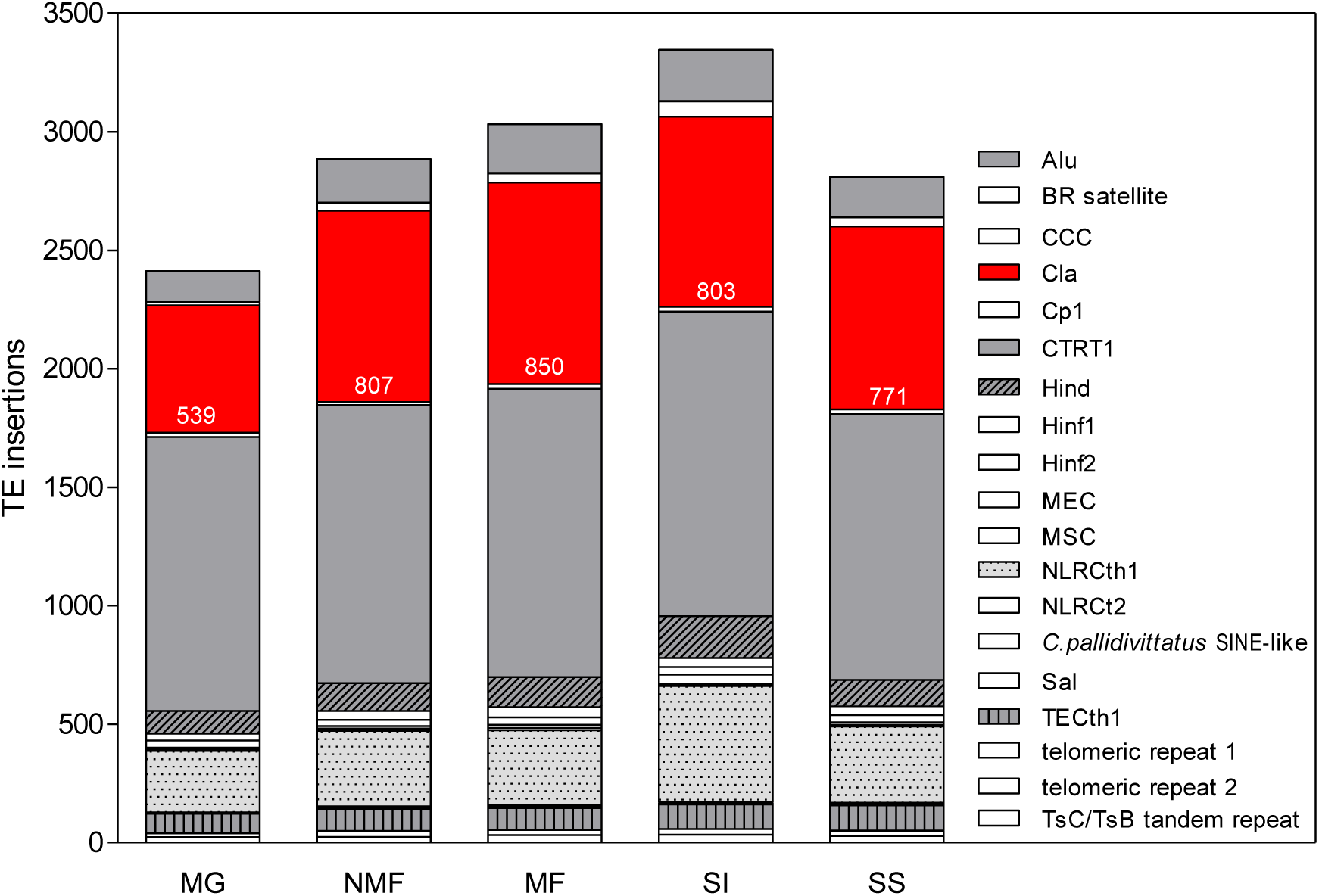
Numbers of detected loci of all transposable element families as represented in the custom TE-library across the whole genome of *Chironomus riparius*. Orientation of the legend refers to the order of the vertical stacked bars. *Cla-element* loci are highlighted in red and numbers of insertion sites are given. All TEs with >100 loci are highlighted in grey and/or pattern.

Among all populations there were only eight TEs inserted close to or into gene regions (Supplementary Table S8), which is significantly less than expected by chance (Supplementary Table S9). Besides insertions into pseudogenes or weak support for the insertion due to low frequencies, only two insertions were localized in predicted genes: the Balbiani Ring repeat was found in the Balbiani Ring protein 2 gene as expected and an Alu element was detected in a retrovirus-related polymerase polyprotein gene (Supplementary Table S8).

### Population genomic analyses of *Cla-element* loci

When analysing the *Cla-element* sequences obtained from the different populations, we did not find a population-specific nucleotide sequence pattern. We randomly selected 100k BLAST hits and aligned them to the *Cla-element* reference sequence of the custom TE-library. The 95% consensus alignment comprised a sequence length of 664 bases containing 430 gaps (data not shown). Thus, variability between *Cla-element* monomers was very high, both on the level of nucleotide substitutions and for insertions and deletions. We did not find any sequence clustering, which would be indicative of *Cla-element* subfamily structure. Due to large variation of homopolymer stretches across the *Cla* sequence, no position of the sequence was invariant throughout the alignment.

Absolute numbers of *Cla-element* insertions varied between populations, ranging from 539 insertions in the Hessian population (MG) to 803 insertions in the Piemonte population (SI, Fig. 1). Based on these numbers of *Cla-element* insertions per population, and the number of reads that were mapped to the *Cla-element* sequence in the TE-library, we estimated the mean *Cla-element* cluster size to be on average 82.8 monomers, ranging from 73.2 (SI) to 89.5 (MF) monomers (Table 1, factor H). The genome-wide number of *Cla* monomer copies was estimated upon the number of mapped reads, ranging between 48,126 in MG and 73,786 in NMF (Table 1, factor I).

In line with the result for all TEs in general, the *Cla-element* was fixed at most insertion-sites (Supplementary Figure S3, disregarding the number of empty sites that arise from comparisons between populations). However, intermediate- and low-frequency insertions were also identified (Supplementary Figure S4) and the mean frequency of segregating sites ranged from 0.57 to 0.64 in the different *C. riparius* populations. We visually validated most of the low-frequency insertions to approve a sufficient coverage and appropriate mapping results in IGV. In all populations, we found low-frequency insertions below 0.25: 16 sites in MG, 16 sites in NMF, 21 sites in MF, 18 sites in SI, and 50 sites in SS. For the two Southern populations (SI – Piemont, SS – Andalusia), *Cla-element* insertions at a frequency of 0.004 were detected in regions with particularly high genome coverage (>1000x) likely resulting from copy number variations.

Low-frequency insertions (here <0.25) might be due to recent transposition activity of the element or purifying selection acting on the element. In order to test for the possibility of ongoing transposition activity, we searched RNA transcriptome data for evidence of *Cla-elements* transcription, as might be expected for active SINE-like retrotransposons. The two 454 transcriptome data-sets from the SRA contained 29 and 27 *Cla-element* sequence reads, most of them present as dimers, none showing traces of polyadenylation or other adjacent mRNA sequences (Supplementary Table S10).

By comparing the *Cla-element* insertion sites between the different populations we obtained presenceabsence patterns. From a total of 1341 *Cla-element* loci, 329 (96 of them fixed) were shared among all populations and are therefore likely ancestral *Cla-element* insertion sites for the species (Supplementary Figure S5).

The relation between population-private and total *Cla-element* loci differed significantly between populations (Supplementary Figure S7). Only 23 private *Cla-element* insertions were identified in Hesse (MG), while the highest number of 118 private insertions was found in the Andalusian population (SS). Relating the number of private *Cla-element* insertions per population to the annual mean temperature of the respective location shows a strong exponential one-phase association (regression r²=0.84, Supplementary Figure S7).

To investigate the structure of the five geographically widespread populations, we estimated the population differentiation based on different genomic elements. On the basis of genome-wide SNPs the mean population differentiation turned out to be very low with the F_ST_ ranging between 0.025 and 0.07, whereas the level of differentiation was at least fourfold higher when considering the mean pairwise F_ST_ on the basis of TEs (0.22 to 0.28). This level of differentiation was even higher when considering the *Cla-element* alone (maximum 0.31 in population pair SI:SS, i.e. Piemont and Andalusia). Concerning levels of differentiation in close vicinity to *Cla-elements* or TEs compared with randomly drawn SNPs from other genome positions, there was no significant difference or visible trend (data not shown). Correlating geographical distance with the mean pairwise F_ST_ revealed that the isolation-by-distance pattern was only significant for SNP-based population differentiation (Pearson one-tailed r=0.57, p=0.0437, Fig. 2B). However, we found a strong correlation between mean population pairwise F_ST_ based on SNPs and TEs (Pearson r=0.87, p=0.0009; only *Cla-element* insertions, Pearson r=0.75, p=0.0125, Fig. 2A).

**Figure 2:**
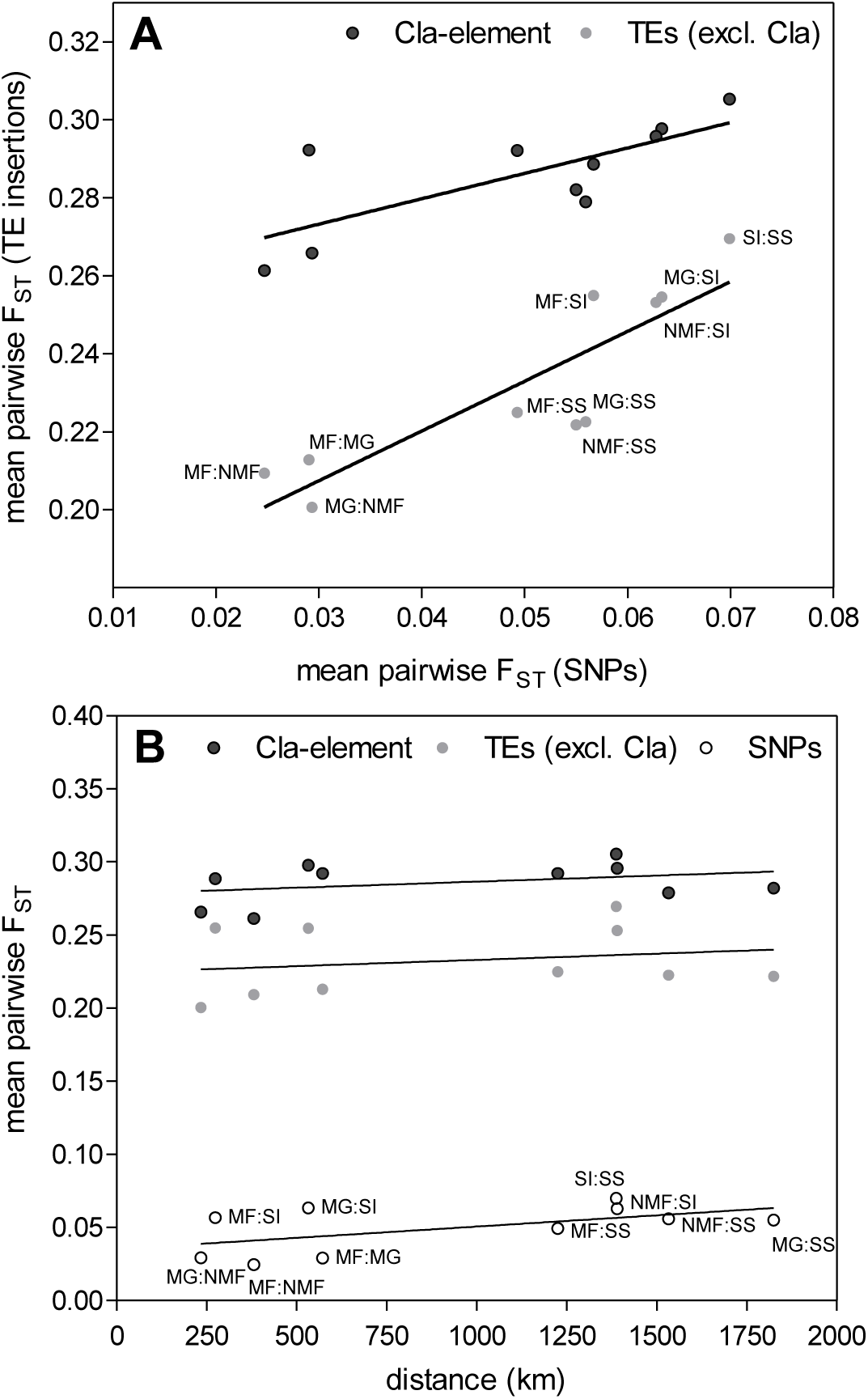
**(A)** Mean pairwise F_ST_s measured with genome-wide SNPs correlated to mean pairwise F_ST_s on the basis of TE insertions (Pearson r=0.8743, p=0.0009) and *Cla-element* insertions (Pearson r=0.7497, p=0.0125). (**B**) Isolation-by-distance pattern as correlation of the geographic distance between the populations with mean pairwise F_ST_s on the basis of genome-wide SNPs (Pearson r=0.5670, p=0.0437), TE insertions (ns), *Cla-element* insertions (ns).

Mean population heterozygosity of private *Cla-element* insertions was significantly lower compared to mean population heterozygosity of private TE insertions excluding *Cla-element*s, and private genome-wide SNPs (Fig. 3). On average, within all populations more than 50% of private *Cla-element* insertions were fixed (Fig. 3 and Supplementary Figure S6). This proportion is significantly higher than the ratio of fixed (relative to all) private TE insertions and genome-wide SNPs (Fig. 3). The fraction of fixed private TE insertions, excluding *Cla-element*s, was on average below 30%, however, still significantly higher than the fixation of genome-wide SNPs private for the different populations (~ 5%).

**Figure 3:**
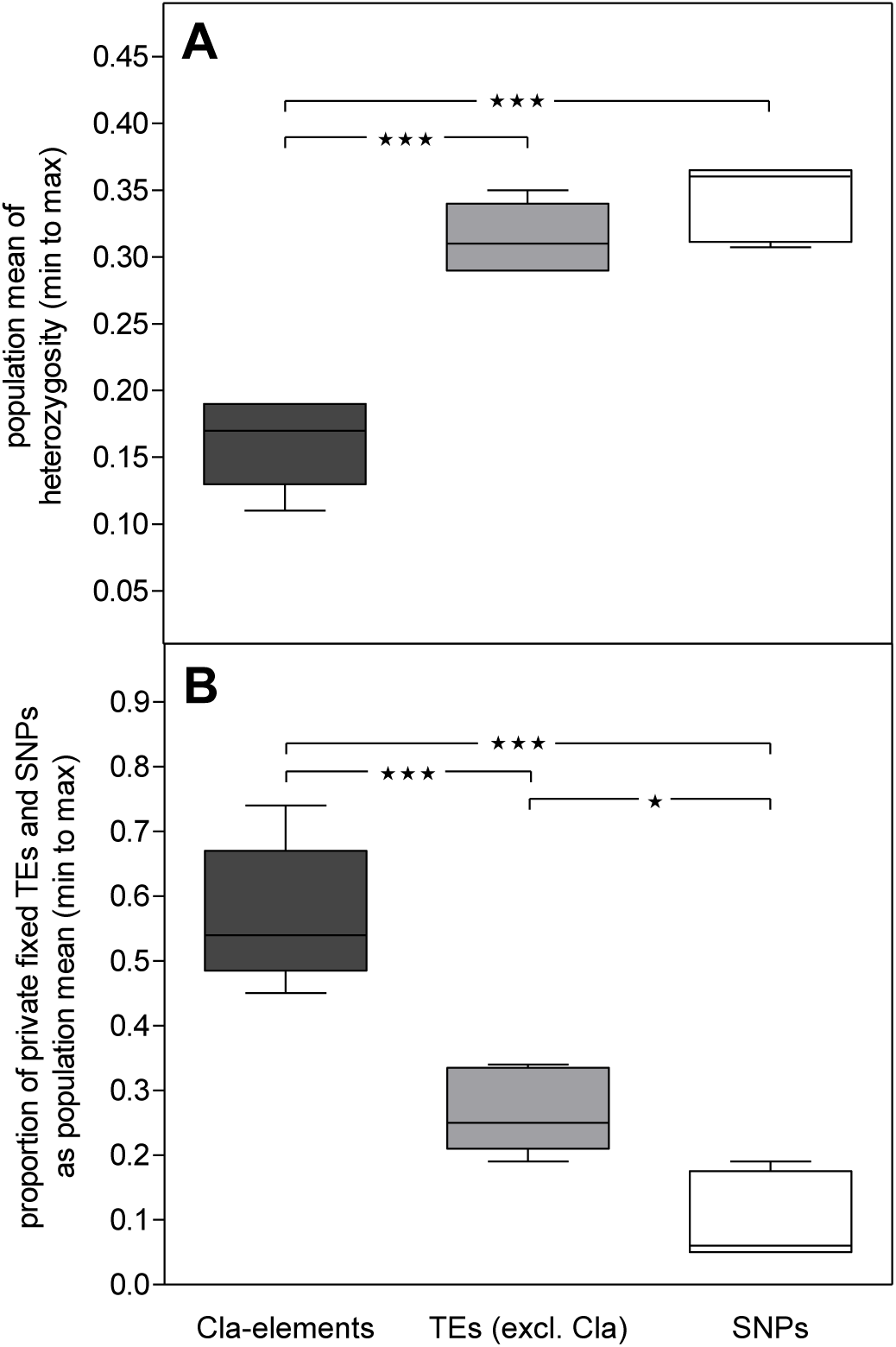
(**A**) Population mean of heterozygosity on private *Cla-element* insertions, TE insertions excluding *Cla-elements*, and genome-wide SNPs, respectively. Heterozygosity is significantly lower for *Cla-element*s compared to TEs and SNPs (one-way ANOVA with Bonferroni post test, significance levels shown as asterisks). (**B**) Population mean of the proportion of private fixed to all private *Cla-element* and TE insertions, and SNPs (one-way ANOVA with Bonferroni post test).

### Chromosomal rearrangements in hybrid individuals

FISH of the *Cla-element* probe was successfully performed for multiple chromosomal sets of each pure and hybrid population (see Supplementary Table S11 for detailed numbers). As expected for *C. riparius*, *Cla-element* bands were distributed across all arms of all chromosomes (Supplementary Figure S2). In order to identify differences in banding patterns between the pure populations, we carefully inspected the chromosomes of hybrid populations. To rule out sample preparation or differential staining artefacts, we required each of our findings to be supported by at least two independent chromosomes. Copy number variations (CNV) inside *Cla-element* clusters were defined as an obvious difference in band width between the two homologous chromosomes. If the *Cla-element* locus was clearly restricted to only one of the homologous chromosomes, we assumed a heterozygous site. In all hybrid populations, we found CNVs or heterozygous *Cla-element* bands that were either homozygous or absent in the pure parental populations, respectively (Fig. 4, marked by asterisks and arrows). Most of these structural differences occurred close to the pericentromeric region of chromosome I or II. Furthermore, partial asynapsis of homologous chromosomes was repeatedly documented on chromosome II in different individuals of the hybrid population MG×SI (Fig 4, circled). Both chromosomal arms were affected by this aberration, which occurred in conjunction with multiple CNVs and heterozygous *Cla-element* bands. A similar effect could be documented on chromosome I of the hybrid population MG×SS, albeit only in a single individual (Fig. 4, bottom).

**Figure 4:**
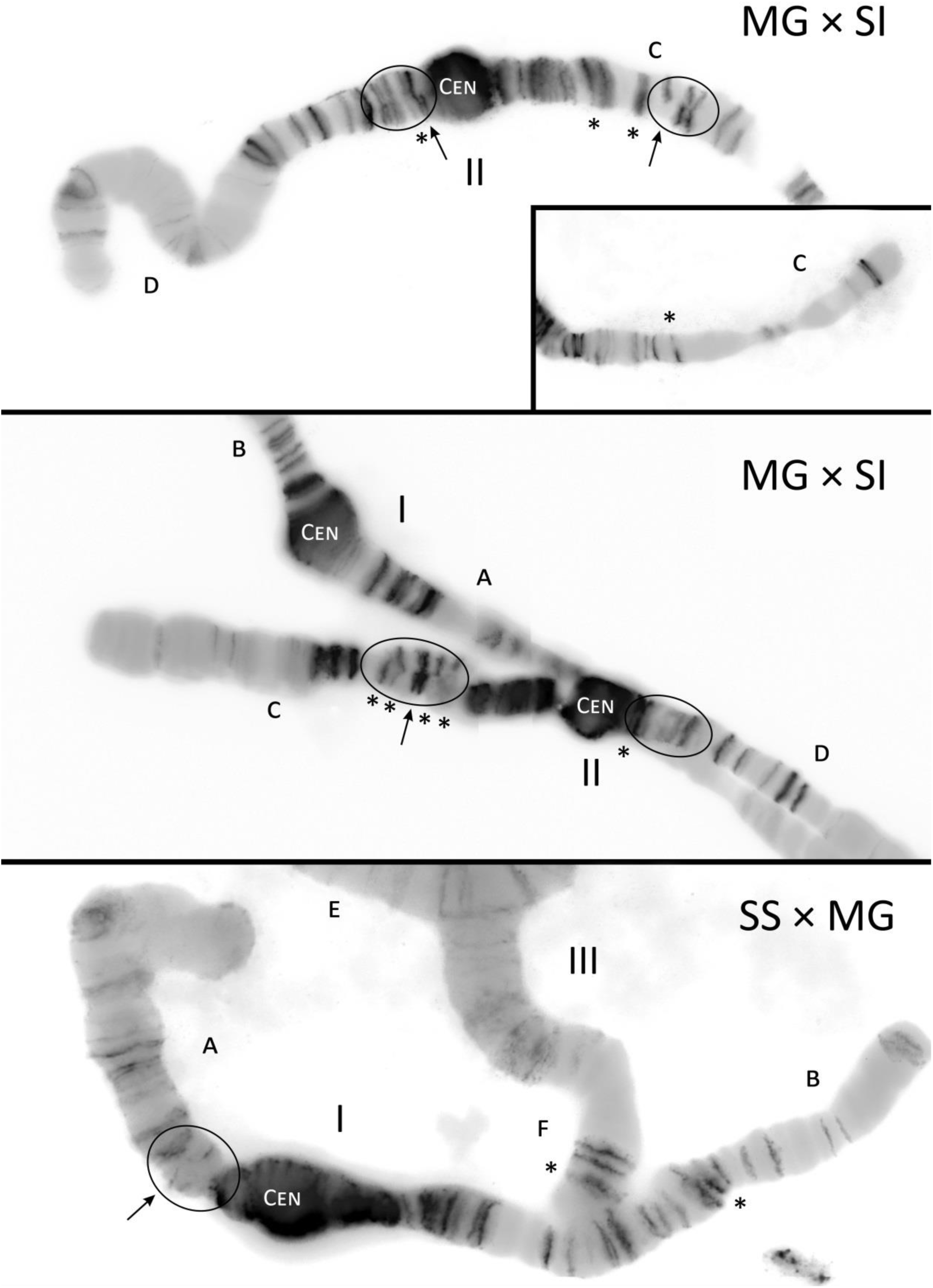
Exemplary fluorescent *in-situ* hybridisation (FISH) micrograph of the *Cla-element* probe hybridized to polytene chromosomes of different *C. riparius* hybrid populations. Pictures of DAPI-stained chromosomes were overlayed by the FISH signal and converted to grey-scale. Copy number variations (asterisks), heterozygous *Cla-element* bands (arrows), and regions of chromosome asynapsis (circles) are highlighted. Chromosomes are designated by Roman numbers (I, II, III), chromosome arms are denoted by letters (A, B, C, D, F), centromeres are labelled (CEN). Graphical insert of chromosomal arm II C (top) results from an additional picture of the identical chromosome.

### Fertility of hybrid populations

The three natural *C. riparius* populations were successfully crossed in both directions resulting in six hybrid populations. Mortality, EmT_50_, PGR, and number of eggs per clutch did not generally differ between pure and hybrid populations (data not shown). The fraction of fertile clutches and number of clutches per female was significantly different in some hybrid populations when compared to the respective pure population (Fig. 5).

**Figure 5:**
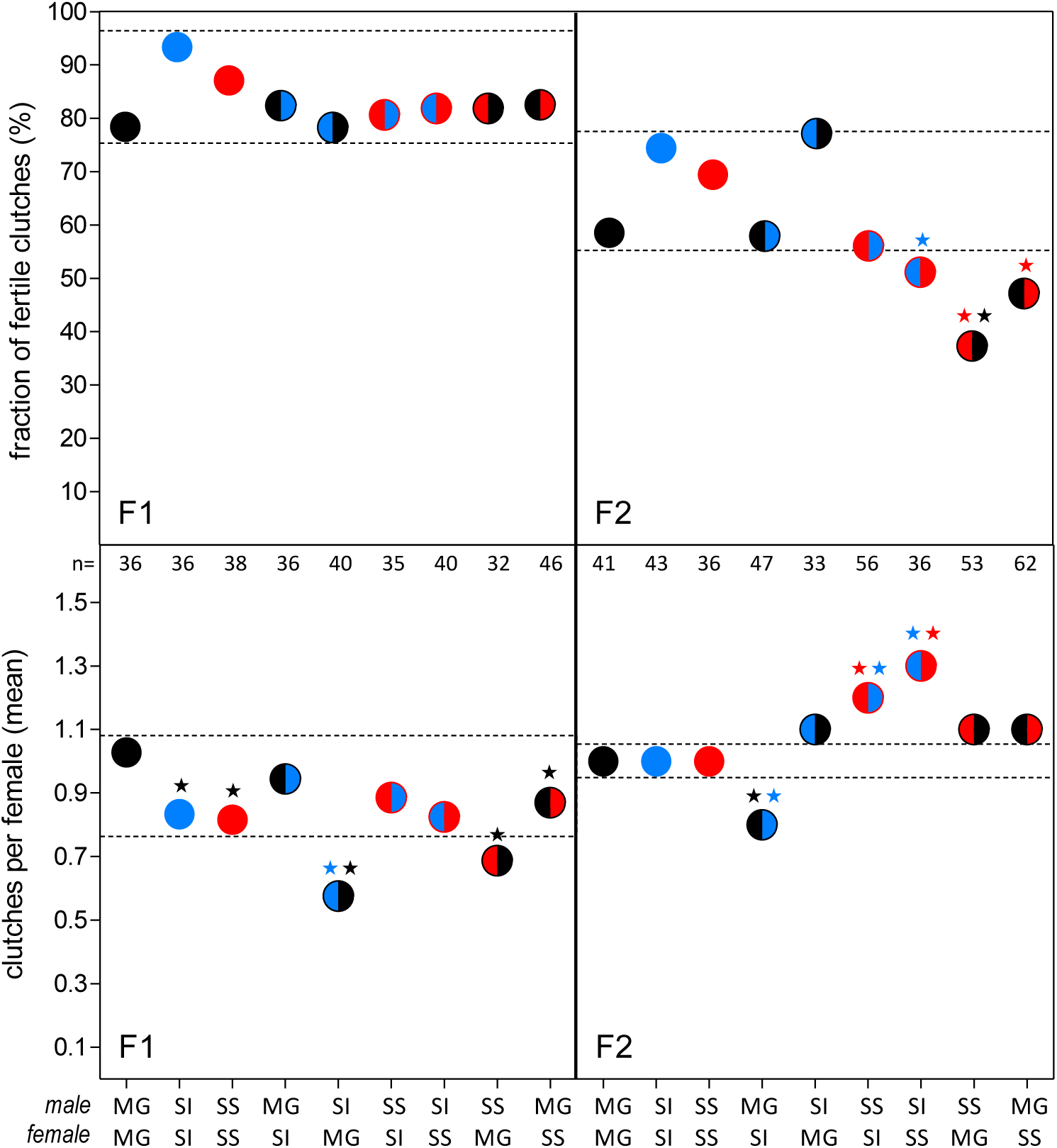
Female fitness parameters in two subsequent generations (F1, left and F2, right) of the hybridisation experiment with different *C. riparius* populations: fraction of fertile clutches (top) and clutches per female (bottom) are indicated. Asterisks mark significant differences to the respective pure population (p<0.05, colours are in accordance to the population code, e.g. black asterisk marks significant difference compared to the black population that is MG, also pure populations were tested against each other; MG - Hessen, SI - Piemonte, SS - Andalucia). Horizontal dashed lines outline the range of variation set by the pure populations.

The fraction of fertile clutches was not affected during the first generation of the hybridisation experiment (Fig. 5 top left). However, fertility of clutches in the following hybrid offspring generation F_2_ of 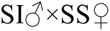 and SS×MG (both crossing directions) decreased significantly compared to at least one of the respective pure populations (Fig. 5 top right). In some hybrid populations the total number of clutches per female was significantly different from the respective pure populations (in both subsequent generations of the experiment; Fig. 5 bottom).

## Discussion

### Draft genome as prerequisite for TE analyses

Computational analyses of genome-wide TE activity require a relatively well assembled and scaffolded draft genome (Ewing 2015). In our draft genome, we could increase the maximum N50 value for genomes of the genus *Chironomus* by the factor of 35 and for all chironomids by the factor of 20 (Supplementary Table S12). In direct comparison with the recently published draft genome of *C. riparius* (Vicoso & Bachtrog 2015), our draft covers a higher fraction of the genome (90% instead of 77.5%), is less fragmented (5,292 instead of 29,677 scaffolds) and has a substantially increased N50 value (272,065 instead of 7,097). From a BUSCO analysis we could assume that the genome is almost complete in terms of its gene set (93 % of core arthropod genes found). The annotation further resulted in a reasonable number of protein-coding genes that is comparable to *C. tentans* from the same genus as well as *Drosophila melanogaster* (15,120 and 13,907 protein-coding genes respectively; Supplementary Table S7; dos Santos *et al.* 2015; Kutsenko *et al.* 2014). The delta of roughly 30 MB of sequence between the presented and the estimated genome size of 200 MB (Schmidt-Ott *et al.* 2009), therefore, is likely mostly due to centromers, telomers, and other large repetitive clusters that cannot be assembled with the existing data.

### Evidence for selection against heterozygous *Cla-elements* as an endogenous genetic barrier among *C. riparius* populations

Using the draft genome and pooled genomic sequencing, we were able to infer the contribution of TEs to population divergence and how they may give rise to endogenous selection barriers between *C. riparius* populations. Despite the large geographic distance between the populations, the population differentiation based on genome-wide SNPs is very low, which is due to a large effective population size and high gene-flow in this species rather than short divergence time (Oppold *et al.* 2016). The population differentiation on the level of *Cla-elements*, and TEs in general, is surprisingly significantly stronger. Many of these loci would have been flagged as potentially under divergent selection in an F_ST_-outlier analysis, indicating that other evolutionary forces are acting on them. However, the significant correlation of mean pairwise F_ST_s measured from genome-wide SNPs with those estimated from all TEs in general and the *Cla-element* insertions in particular (Fig. 2A) suggests on the other hand that drift nevertheless shaped the overall population differentiation pattern of these genomic elements. Many private insertions indicate that either ongoing TE activity after population-split or purifying selection by differential purging (Charlesworth & Charlesworth 1983) played a major role in TE population dynamics (reviewed in Barron *et al.* 2014). We therefore focussed on private TE and *Cla-element* insertions compared to private SNPs. For the majority of the latter, we can reasonably assume that they occurred after the respective population splits. If neutral, the vast majority of private markers should therefore follow the same population genetic trajectories, governed by the respective demographic population parameters. Given the low overall differentiation (mean F_ST(SNPs)_= 0.05), indicative of high inter-population gene-flow (Slatkin 1987), we would expect strictly neutral private TE insertions to segregate within populations at about the level of private neutral SNPs. This is not the case, as shown by the significant difference in heterozygosity and the higher proportion of fixed private TEs, in particular for the *Cla-elements* (Fig. 3, Supplementary Figure S6). Non-neutral, i.e. endogenous selective processes acting on TEs in general and on the *Cla-elements* in particular are therefore necessary to explain the observed patterns. We suggest that the heterozygous state of tandem-repetitive *Cla-element* clusters as such in an individual may be deleterious. The following scenario may explain our findings: To explain the presence of fixed private insertions, there has to be a mechanism through which new insertions can initially escape negative selection. It can reasonably be assumed that short *Cla-element* monomers are nearly neutral and may, thus, by chance drift to frequencies at which homozygous insertions occur. The latter is necessary to start expansion in a tandem-like fashion by unequal crossing-over. This process finally enhances the formation of large tandem-repetitive clusters because generally the frequency of unequal crossing-over events or slipped strand mispairing increases with the length of an allele (Levinson & Gutman 1987; Levinson *et al.* 1985). Depending on their population frequency, such clusters might either become fixed or eliminated by selection. We are, however, not able to test this hypothesis as our data does not allow for length comparisons of *Cla-element* clusters. As potential deleterious mechanisms, secondary structures formed by tandem-repetitive *Cla-element* clusters due to hairpin loop formation and the marked inherent DNA-curvature (Hankeln 1990; Israelewski 1983; Schmidt 1984) might lead to complex epistatic interaction triggering DMI (Brown & O'Neill 2010) and ectopic recombination events, as observed for classical TEs in *Drosophila* (Barron *et al.* 2014; Petrov *et al.* 2011). Suggestively, chromosomal regions containing *Cla-element* clusters are associated with known chromosome breakpoints in *C. riparius* (Bovero *et al.* 2002), and also in our study chromosomal pairing aberrations were observed in conjunction with heterozygous *Cla* clusters (Fig. 4). Ectopic recombination events are more likely for heterozygous TEs and their deleterious consequences should be under strong negative selection (Montgomery *et al.* 1991). As this inevitably leads to differential fixation of TE insertions among populations, in particular inter-population hybrids during the first few generations are expected to harbour an increased amount of heterozygous insertion sites and, thus, experience the consequences of multiple DMI as an endogenous selection pressure, as observed here. In principle, adjacent variation should hitchhike during fixation of a *Cla-element* and consequently show a similar differentiation pattern. However, this is not the case, suggesting that most instances are relatively old events where recombination and mutation have already erased any potentially initially present signs of hitchhiking during the fixation process, as predicted by Bierne *et al.* (2011). In general, the abundance of TEs in the *C. riparius* genome reflects the expected distribution pattern of significantly less TEs inserted into protein-coding regions than expected by chance, supporting their deleterious influence on conserved coding regions (Pasyukova *et al.* 2004).

We here propose a TE-based mechanism for the induction of DMI as basis for endogenous selection. Thus, this phenomenon does not seem to only arise from classical TEs, but also from transposable tandem-repetitive minisatellite-like sequences with a presumably passive mode of transposition. The mechanism proceeds aside from the concept of genomic conflict (Crespi & Nosil 2013) and has, to our best knowledge, so far not been empirically investigated.

### Post-population split transpositional activity of the *Cla-element* in *C. riparius*

To contribute to population divergence, TEs must have either been active after the population split or TEs segregating at the time of the population split must have been fixed differentially thereafter. Several aspects of our data point towards a post-split transposition activity of the *Cla-element* in the analysed *C. riparius* populations. First, there are many population-specific *Cla-element* insertion sites resulting in a divergent genomic TE pattern among populations (Fig. 1). A scenario of differential fixation is only imaginable if the pre-split *Cla-element* was subject to different selective forces than today. If the same negative selection pressure inferred to act on heterozygous *Cla-element* insertions in the current populations applied back then, most of the *Cla-element* insertions would have been fixed as they are today and could not get lost differentially. Sequence comparisons of the *Cla* consensus sequence revealed high similarity to the sister species *C. piger* (3.7% *Cla-element* divergence, (Hankeln & Schmidt 1987) and even to the more distantly related *C. luridus* (Ross *et al.* 1997), hence, it is unlikely that structural features of the element on which selection may act, were any different in the ancestral *C. riparius* population. We therefore suggest that the *Cla-element* has at least been active after the establishment of the different populations at their respective geographical site. Regarding the high variability of the *Cla* sequence identified via the BLAST analysis, it might be argued that the *Cla-element* has generally accumulated too many mutations to maintain activity until today. However, classical TEs such as the ETn elements in the mouse genome are transposed from a few master copies of intact feature architecture that will ensure continued active transposition (Ribet *et al.* 2004). With the applied method we cannot distinguish sequence variability in regard of the respective insertion site, and thus no statement about the *Cla* sequence at population-private or non-fixed sites can be made. Furthermore, the transposition mechanism of the *Cla-element* is still unknown, and so are the necessary sequence features for activity maintenance.

Second, the existence of insertion sites still segregating in the population indicates that these sites have been introduced, either by *de novo* insertion or migration, recently enough to avoid fixation by drift. An active TE will constantly create new insertions that will start at low frequencies in the population and that are exclusive to this population. Only about 25% of all *Cla-element* loci were shared among all populations and are therefore likely to belong to the most ancient species-specific insertions (Supplementary Figure S5). The detected population-specific low frequency (<0.25) *Cla-element* insertions can be best explained with active transposition of the element after the population split. However, negative selection against segregating (i.e. heterozygous) *Cla-element* insertions could also be the reason for their relatively low amount.

In the two Southern-most populations (SI - Piemonte, SS – Andalucia), we detected insertions at very low frequencies (0.003 – 0.01) in different genomic regions with high coverage, respectively. On the one hand, this might be an artefact due to *Cla-element* CNV. On the other hand, these regions might be heterochromatic with a high content of so far unknown repetitive elements that are not well assembled and not masked in the draft genome, thus explaining the high coverage. Such regions being presumably neutral, they allow for new insertions to easily accumulate there (Cridland *et al.* 2013). Therefore, these low frequency *Cla-element* insertions might be traces of recent transposition events in these two populations. The higher proportion of private *Cla-element* insertions and low-frequency insertions in the Southern populations (SI – Piemonte, SS – Andalucia) suggest that the *Cla-element* is more active there. Consistently, significantly less population-private *Cla-element* insertions were found in Northern populations (MG – Hessen, NMF – Lorraine). Additionally, this pattern is positively correlated (exponential one-phase association) to the annual mean temperature at the respective location (Supplementary Figure S7). These results fit very well to a recent study showing that the evolutionary “speed” of the very same *C. riparius* populations depends on the temperature dependent number of generations per year and the possibly also increased mutation rate (Oppold *et al.* 2016). It is conceivable that the observed quantitative differences in population TE content are also temperature related.

Third, the presence of monomeric or dimeric *Cla-element* transcripts in *C. riparius*, as evidenced by transcriptome data (Supplementary Table S10), Northern hybridization (Hankeln & Schmidt 1987), and RT-PCR (Martinez-Guitarte *et al.* 2012) further hints at an ongoing *Cla-element* transposition, presumably *via* RNA intermediates that are then reverse transcribed and re-integrated into the genome. Monomeric *Cla-elements* are structurally similar to classical SINE retroelements (Hankeln & Schmidt 1987). However, SINEs depend on active LINEs (long interspersed nuclear elements) for their retrotransposition machinery (Wicker *et al.* 2007). In fact, there is indirect evidence for ongoing transpositional activity of at least one LINE in *C. riparius* (NLRCth1; Zampicinini *et al.* 2011) that could provide the necessary enzymatic components for passive *Cla-element* transposition. It remains unclear, however, if *Cla-elements* predominantly retrotranspose as mono- or oligomers. Most *Cla* sequences are organized as minisatellite-like, interspersed clusters, which might also use DNA-based transposition, e.g. by excision and re-integration of extrachromosomal copies (reviewed in Kazazian 2004).

Taken together, population-specific *Cla-element* abundance, *Cla-element* insertions at low and intermediate frequencies in the respective populations, and the presence of *Cla* transcripts strongly suggests recent transpositional activity of the element in the investigated populations. The effect of drift or purifying selection acting on *Cla-element* insertions could alternatively explain the divergent abundance and segregating insertion sites, but not the *Cla* transcripts.

### Incompatibilities on the phenotypic level of chromosomes among *C. riparius* populations

In-situ hybridisation of *Cla-elements* to polytene chromosomes of the different pure and hybrid populations showed an additive *Cla-element* pattern in F1 hybrid individuals, as expected. A drastic burst of *Cla-element* activity creating new loci during early embryo development was therefore not observed, in contrast to the activation of the P-element in hybrid dysgenesis of *D. melanogaster* when only one strain is carrying the P-element (Rio 1990). The additive effect of the *Cla-element* pattern in hybrid individuals led to several heterozygous sites, sometimes showing partial asynapsis of the homologous chromosomes in this region (Fig. 4). Hybridising *C. riparius* with *C. piger*, the sister species that is free of *Cla-element* insertions across chromosomal arms, results in very strong asynapsis of homologous chromosomes (Hägele 1984; Schmidt 1984). Based on these findings and the correlation of *Cla-element* distribution with sites of chromosomal rearrangements and breakage (Bovero *et al.* 2002), we propose that the heterozygous *Cla-element* pattern leads to incorrect synapsis of homologous chromosomes as an obvious evidence of DMI on the level of chromosome structure.

### Crossing of *C. riparius* populations affects hybrid fitness

Whether the detected chromosome aberrations might actually directly influence the fitness of individuals is difficult to test with our data. Our observation of reduced fertility of hybrid clutches (Fig. 5) is in line with the observations of reduced egg hatch in hybrids of different *C. riparius* strains that suffered from the HLE dysgenesis syndrome (Hägele 1995). Impairment of clutch fertility was especially prominent in crosses of the Hessen population (MG) with Andalusian population (SS), suggesting a population-specific dysgenesis potential (Hägele 1995). The hybrid crosses between MG × SS and SI × SS show reduced clutch fertility as well as an increased number of clutches per female potentially as a compensatory effect. Future experiments on different phenotypic traits will be necessary to investigate if hybrid individuals show intermediate phenotypes and reduced fitness in regard to the respective environmental conditions of the natural pure populations, e.g. temperature regimes. However, as an intrinsic genomic turnover component (Dover 1982), TEs drive population divergence independently of the local environment (Bierne *et al.* 2011). If this results in negative fitness effects for hybrids, they are not necessarily expected to show intermediate phenotypes between clines. For the present study, it could so far be documented that fertility of offspring is significantly reduced when populations of differing genetic backgrounds are hybridised, which eventually might enhance RI. In *Drosophila*, the dysgenesis syndromes involving the P-, I- or the *hobo* element are known to be restricted to one crossing direction (transposon-carrying males × transposon-free females, Bingham *et al.* 1982; Bucheton *et al.* 1984; Galindo *et al.* 1995). Lack of such a bias in our study indicates that the underlying mechanism may be different in *C. riparius* and can be better explained with the general concept of DMI that irrespective of specific loci can result in endogenous genetic barriers.

## Conclusion

We could show that TEs might be an underestimated source of genomic differentiation and hence; the divergence; and ultimately perhaps even speciation of conspecific populations. This has to be accounted for when investigating the genomic background of possible local adaptation of populations as already suggested by Bierne *et al.* (2011). The improved and annotated *C. riparius* draft genome will offer a valuable genomic resource beyond the focus of this study.

## Acknowledgements

We thank Steffen Lemke from the Centre for Organismal Studies Heidelberg, providing initial support for the genome assembly. The 1KITE project (www.1kite.org/), namely Karen Meusemann, gratefully provided us with additional transcriptome data for the genome annotation process. We also thank Ingo Ebersberger for advice and fruitful discussions, as well as Daniel Barbash for helpful comments. Funding of the project was provided by DFG (PF390/7-1; Ha2103/7-1). Parts of this research were conducted using the supercomputer Mogon and advisory services offered by Johannes Gutenberg University Mainz (www.hpc.uni-mainz.de), which is a member of the AHRP and the Gauss Alliance e.V. T.H. gratefully acknowledges support by the Centre for Computational Sciences, Johannes Gutenberg University Mainz (CSM/SRFN). Robert Kofler acknowledges funding within the ERC Grant “ArchAdapt”.

## Data accessibility

Genome sequencing raw reads are deposited in the EMBL-EBI European Nucleotide Archive (accession numbers pending). Re-sequencing raw reads are deposited in the NCBI Short Read Archive (accession numbers pending). Transcriptome sequencing raw reads are deposited in the EMBL-EBI European Nucleotide Archive (accession numbers pending). The final DNA sequence assembly is deposited in the EMBL-EBI European Nucleotide Archive (accession number pending). The genome annotation file is available from. Mapping data in *bam format is deposited at Dryad (accession numbers pending). Coordinates of sampling locations are available in the Supplementary Material.

## Authors’ contribution

A.-M.O., T.H. and M.P. conceived the study; T.H., F.R., and U.S.-O. performed genome sequencing; H.S, S.L.H., and F.R. assembled the draft genome; H.S., F.D., and S.L.H. annotated the draft genome; A.-M.O. performed sequencing of population Pool-Seq genome scans, A.-M.O., R.K., and M.P. performed population genomic analyses, M.R. and A.-M.O. performed the hybridisation experiment, M.R., B.W., T.H., and E.S. performed and analysed FISH of polytene chromosomes, A.-M.O., H.S., S.L.H., and M.P. drafted the manuscript.

